# MaxQuant and MSstats in Galaxy enable reproducible cloud-based analysis of quantitative proteomics experiments for everyone

**DOI:** 10.1101/2022.01.20.477129

**Authors:** Niko Pinter, Damian Glätzer, Matthias Fahrner, Klemens Fröhlich, James Johnson, Björn Andreas Grüning, Bettina Warscheid, Friedel Drepper, Oliver Schilling, Melanie Christine Föll

## Abstract

Quantitative mass spectrometry-based proteomics has become a high-throughput technology for the identification and quantification of thousands of proteins in complex biological samples. Two de facto standard tools, MaxQuant and MSstats, allow for the analysis of raw data and finding proteins with differential abundance between conditions of interest. To enable accessible and reproducible quantitative proteomics analyses in a cloud environment, we have integrated MaxQuant (including TMTpro 16/18plex), Proteomics Quality Control (PTXQC), MSstats and MSstatsTMT into the open-source Galaxy framework. This enables the web-based analysis of label-free and isobaric labeling proteomics experiments via Galaxy’s graphical user interface on
public clouds. MaxQuant and MSstats in Galaxy can be applied in conjunction with thousands of existing Galaxy tools and integrated into standardized, sharable workflows. Galaxy tracks all metadata and intermediate results in analysis histories, which can be shared privately for collaborations or publicly, allowing full reproducibility and transparency of published analysis. To further increase accessibility, we provide detailed hands-on training materials. The integration of MaxQuant and MSstats into the Galaxy framework enables their usage in a reproducible way on accessible large computational infrastructures, hence realizing the foundation for high throughput proteomics data science for everyone.

## INTRODUCTION

Mass spectrometry-based proteomics is a standard technique for the identification and relative quantification of thousands of proteins in complex samples. A common aim is to identify proteins that are differentially abundant between conditions of interest. Two standard software tools for data dependent acquisition (DDA)-based quantitative proteomics are MaxQuant ^1,2^ and MSstats ^3,4^. Together they allow for a typical quantitative shotgun proteomics analysis workflow. MaxQuant is a standalone freeware that takes raw data as input and performs protein identification and quantification. MaxQuant supports all common protein quantification methods such as label-free, label-based and isobaric labeling ^1,5^. MSstats is a Bioconductor R package for finding proteins that are differentially abundant in different conditions. It uses flexible linear models to analyze label-free proteomics experiments with complex designs ^3^. Recently, MSstatsTMT was released for the statistical modeling of isobaric labeling quantification data e.g. iTRAQ (isobaric tags for relative and absolute quantitation) or TMT (tandem mass tag) ^4^.

Typically, a quantitative proteomics analysis requires several steps: First, all software needs to be installed. Often this is done on a shared lab workstation with sufficient computational power. Next, the MaxQuant run is started and once it is finished the results may be inspected manually or with a dedicated software such as the PTXQC R package to obtain a direct quality control report ^6^. Afterwards, the MaxQuant result files are loaded into the R programming environment for processing and statistical analysis in MSstats. In contrast to many other proteomics software, MaxQuant and MSstats are compatible with powerful computational infrastructures such as high-performance computing (HPC) systems and cloud environments ^7,8^. This is required as technical advancements in sensitivity, mass resolution and acquisition speed lead to larger file sizes and increasing number of samples per experiment ^7^. With the steadily expanding availability of instrumentation, proteomics experiments are increasingly widespread and complex. This emphasizes the need for easily accessible and scalable software solutions. However, even for HPC and cloud compatible software, monetary hurdles and technical complexity of software installation and maintenance severely hamper the access to high throughput analysis ^7^. Reproducible, and thus trustable high-throughput analyses require even more computational skills to control software versions and dependencies ^7^. Even the most detailed methods section cannot ensure reproducible analyses if not every researcher has access to the same software, software version, computing environment and computational resources ^9^.

Here, we present the integration of MaxQuant and MSstats into the Galaxy framework to enable accessible and reproducible quantitative proteomics analyses in a cloud environment. The biomedical data analysis platform Galaxy is an open-source, free-to-use web-based service with a graphical user interface that can schedule pre-installed tools on large compute resources and public clouds while recording all provenance data from parameters to the tool version and information on analysis workflows ^10^. It allows sharing of complete analysis histories and workflows and therefore provides a platform on which high throughput analyses can be executed and repeated by everyone. Galaxy already offers thousands of tools for many different omics domains, including a variety of tools for explorative proteomics, such as msconvert ^11^, SearchGUI ^12^, PeptideShaker ^13^, OpenMS ^14^, OpenSwath ^15^, and DIAumpire ^16^. Thus, MaxQuant and MSstats in Galaxy not only enable classical DDA-based quantitative proteomics analyses but may also be integrated with other Galaxy tools into standardized, shareable workflows. With the integration of MaxQuant and MSstats into the Galaxy framework, we enable every researcher to run quantitative proteomics analysis of a quasi unlimited number of files on public cloud infrastructures.

## IMPLEMENTATION

### Technical details of the MaxQuant and MSstats integration into Galaxy

We have integrated MaxQuant (versions 1.6.3.4, 1.6.10.43, 1.6.17.0), PTXQC (versions 0.92.6, 1.0.9, 1.0.10), MSstats (versions 3.20.1, 3.22.0) and MSstatsTMT (versions 1.8.0, 1.8.2) functionalities into the Galaxy framework. First, we have built Bioconda recipes and BioContainers that manage all the dependencies for MaxQuant, PTXQC, MSstats and MSstatsTMT respectively ^17,18^. These allow for easy software installation while having full version control in any (Linux-based) environment. Therefore, they are also beneficial for applications outside of the Galaxy framework. Within the Galaxy framework, Bioconda recipes and Biocontainers allow installation of multiple versions of the same tool and easy switching between them, which allows full reproducibility even of older analyses. Lastly, we built so-called Galaxy wrappers that define the input parameters in Galaxy’s graphical user interface and link them to the software executables.

In this way, four new Galaxy tools were built: ‘MaxQuant’ including PTXQC functionality, ‘MaxQuant (using mqpar.xml)’ including PTXQC functionality, ‘MSstats’ and ‘MSstatsTMT’.

### MaxQuant in Galaxy

Two MaxQuant tools were integrated into Galaxy framework: One uses the mqpar.xml parameter file as input while the other allows setting parameters directly in the tool user interface. Both MaxQuant tools offer the same options for raw data, database input files and output files. Raw data is accepted in the Thermo RAW file format as well as in the open standard formats mzXML and mzML ^19,20^, which can be obtained by converting any vendor-specific RAW format with the msconvert software ^11^ that is also available in Galaxy. Single or multiple FASTA files are allowed as database input and the ‘parse rules’ can directly be adjusted in the user interface and do not require an additional configuration step. All common MaxQuant files are offered as output options. The PTXQC R script is directly integrated into the MaxQuant tools and allows the optional creation of a QC report following the MaxQuant run.

The ‘MaxQuant (using mqpar.xml)’ tool runs MaxQuant with the input parameters specified in a mqpar.xml parameter file that was created beforehand, e.g. by using the traditional MaxQuant software. The intended use-case is to scale from a local installation easily to large compute resources using Galaxy. In addition to the selection of input files, the only parameters that have to be set are the “parse rules” for the FASTA file, the PTXQC parameters and the selection of output files.

Since mqpar.xml files do not always exist or might need complicated adjustments, we have built an additional ‘MaxQuant’ tool that allows specifying the most crucial parameters directly in the Galaxy user interface. The tool is separated into five categories: Input options, Search options, Protein quantification, Parameter group and Output options. In contrast to the original MaxQuant software, the experimental setup, which includes file name, experiment name, fraction and post translational modifications needs to be specified in a tab-separated values file outside Galaxy. Custom modifications cannot be configured by the user. They need to be added by a Galaxy tool developer, but once the modifications are installed they will remain in all following tool versions. We have integrated the modifications for TMTpro-16plex and TMTpro-18plex, allowing the user to use these quantification options directly without any additional installation steps. Inside the Galaxy tool, the specified parameters are transferred via an additional python script into the mqpar.xml parameter file, which is then used to launch MaxQuant.

### MSstats in Galaxy

Two MSstats Galaxy tools were built based on the Bioconductor R packages MSstats and MSstatsTMT, which analyze quantitative proteomics data from label-free and isobaric labeling data, respectively. The MSstats and MSstatsTMT Galaxy tools cover the entire statistical analysis workflow from importing and converting results from quantitative proteomics software to protein summarization, protein quantification, and group comparison. For this workflow, the full set of parameters is adjustable via the Galaxy tool interface and MSstats converter for MaxQuant, OpenSwath (only MSstats), OpenMS and Proteome Discoverer (only MSstatsTMT) are included. Like in the original software, an additional file is needed to specify experimental annotations such as condition, biological and technical replicates. Quantitative proteomics data from not supported software such as Skyline and Progenesis can be converted and annotated outside Galaxy, for example in a text editor, into the MSstats specific table format.

The desired comparison between conditions requires an additional tab-separated value file that defines the comparison matrix. For each analysis step, the user can select the result tables and visualizations of interest.

## RESULTS AND DISCUSSION

### Access to MaxQuant and MSstats in Galaxy

To enable every researcher to perform reproducible and scalable quantitative proteomics analyses, we have integrated two de-facto standard tools, MaxQuant and MSstats, into the Galaxy framework. According to the modular tool structure in the Galaxy framework, we have built four new tools: ‘MaxQuant’ including PTXQC functionality, ‘MaxQuant (using mqpar.xml)’ including PTXQC functionality, ‘MSstats’ and ‘MSstatsTMT’. These tools are available via the Galaxy toolshed ^25^, which is the central tool repository from which Galaxy administrators can install the tool on any Galaxy server, including the more than 125 public Galaxy servers. The described tools are already installed on several public Galaxy servers, where everyone can create a free user account and use the graphical user interface to adjust tool parameters and run the tools on public computing infrastructure. Only internet access and a web-browser are needed to access these public Galaxy instances. In the case of the European Galaxy server (https://usegalaxy.eu), thousands of cores and dozens of terabytes of RAM are available (de.NBI cloud), allowing for the comprehensive analysis of large proteomics datasets (Figure 3). Table S1 provides links to the tools in the Galaxy toolshed and on the European Galaxy server.

### Quantitative proteomics in the Galaxy framework

In combination, MaxQuant and MSstats enable protein quantification and differential abundance analysis of label-free, TMT and iTRAQ proteomics data. Within Galaxy, MaxQuant and MSstats can be operated individually or together in a workflow. Workflows require the user to only start the analysis once because the generation of the MaxQuant results automatically triggers MSstats to continue with the analysis. Regardless if the analysis is performed step by step or via workflows, Galaxy histories are generated. A history contains all intermediate and result files together with all metadata required for transparency and reproducibility such as tool name and tool versions and the used parameters and input files. Histories and workflows can be either shared privately with collaborators or publicly for example as part of peer-reviewed publications.

Several hundred to thousands of Galaxy tools are pre-installed on every Galaxy server and enable high levels of interoperability. Therefore, MaxQuant and MSstats Galaxy tools seamlessly integrate into the already existing tool landscape of proteomics ^26–28^, metabolomics ^29–31^ and many more omics disciplines that allow complex, large-scale proteomics and multi-omics analysis ^32^ in a reproducible manner (Figure 2). MSstats is the first Galaxy tool specialized for statistical analysis of quantitative proteomics data. It is not only compatible with MaxQuant but also other proteomics software that is available in Galaxy such as OpenMS and OpenSwath ^33^ and therefore expands analysis options inside Galaxy. The tab-separated values file outputs of MaxQuant and MSstats are compatible with the many text manipulation tools in Galaxy that allow for example filtering, sorting, computing, summarizing and visualization. All tab-separated values files are furthermore compatible with other downstream Galaxy tools such as protein annotation, Gene Ontology (GO) annotation and enrichment analysis.

**Figure 1:**
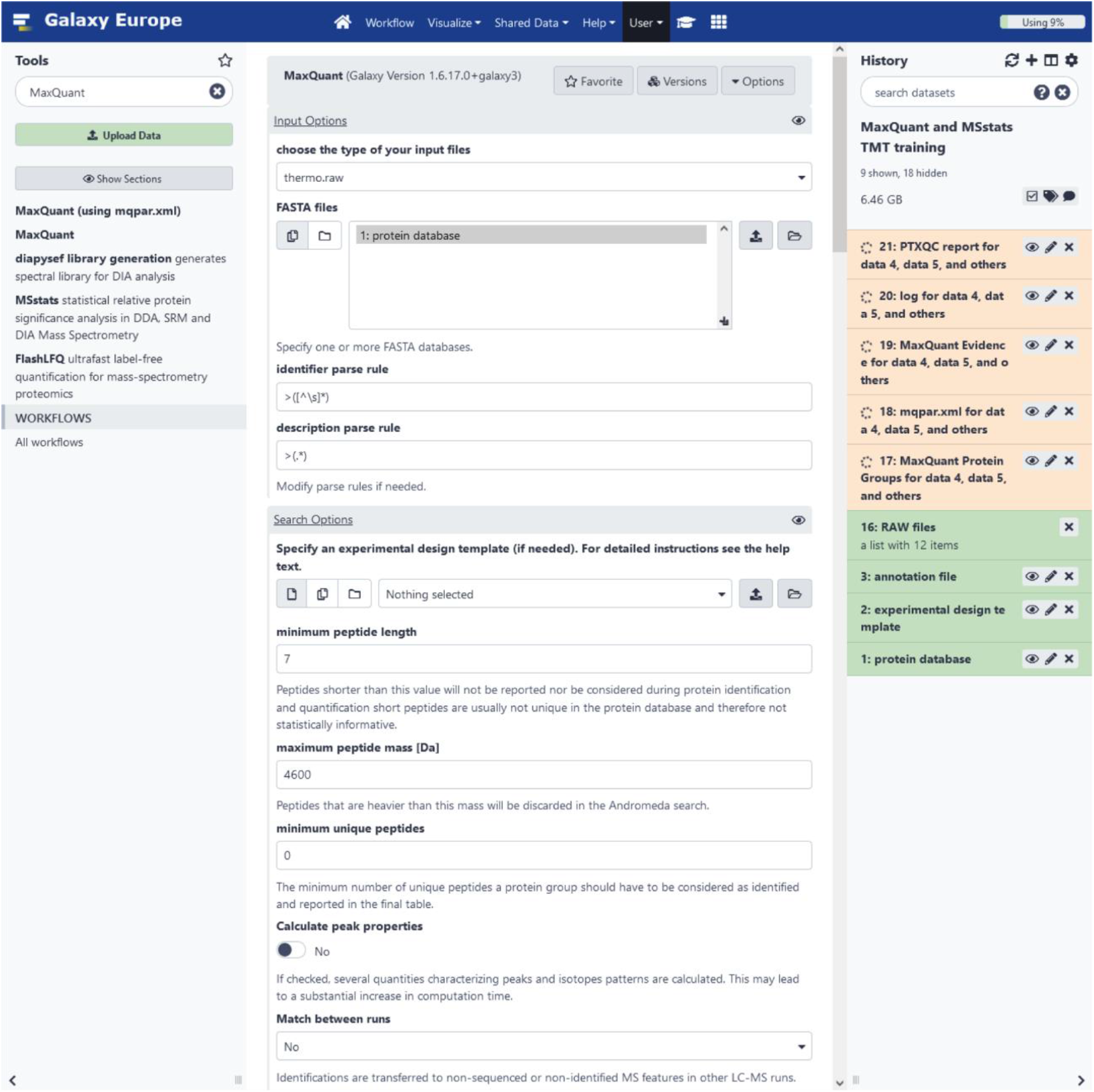
The Galaxy user interface with the MaxQuant tool. The Galaxy user interface is separated into three panels. The tool box on the left, the main panel in the middle and the history on the right side. Tools can be searched by categories or text search in the tool box. The selected tool will be opened in the main panel. Here, the top section of the MaxQuant tool in Galaxy is shown. Files from the history can be selected as input files and parameters for the MaxQuant run can be adjusted. After starting the tool, its output files are sent to the history. The history panel shown contains the protein database, experimental design template, annotation file for MSstats and RAW files. The yellow entries in the history are the output files of a MaxQuant run in progress. Once finished, the entries will turn green.

**Figure 2:**
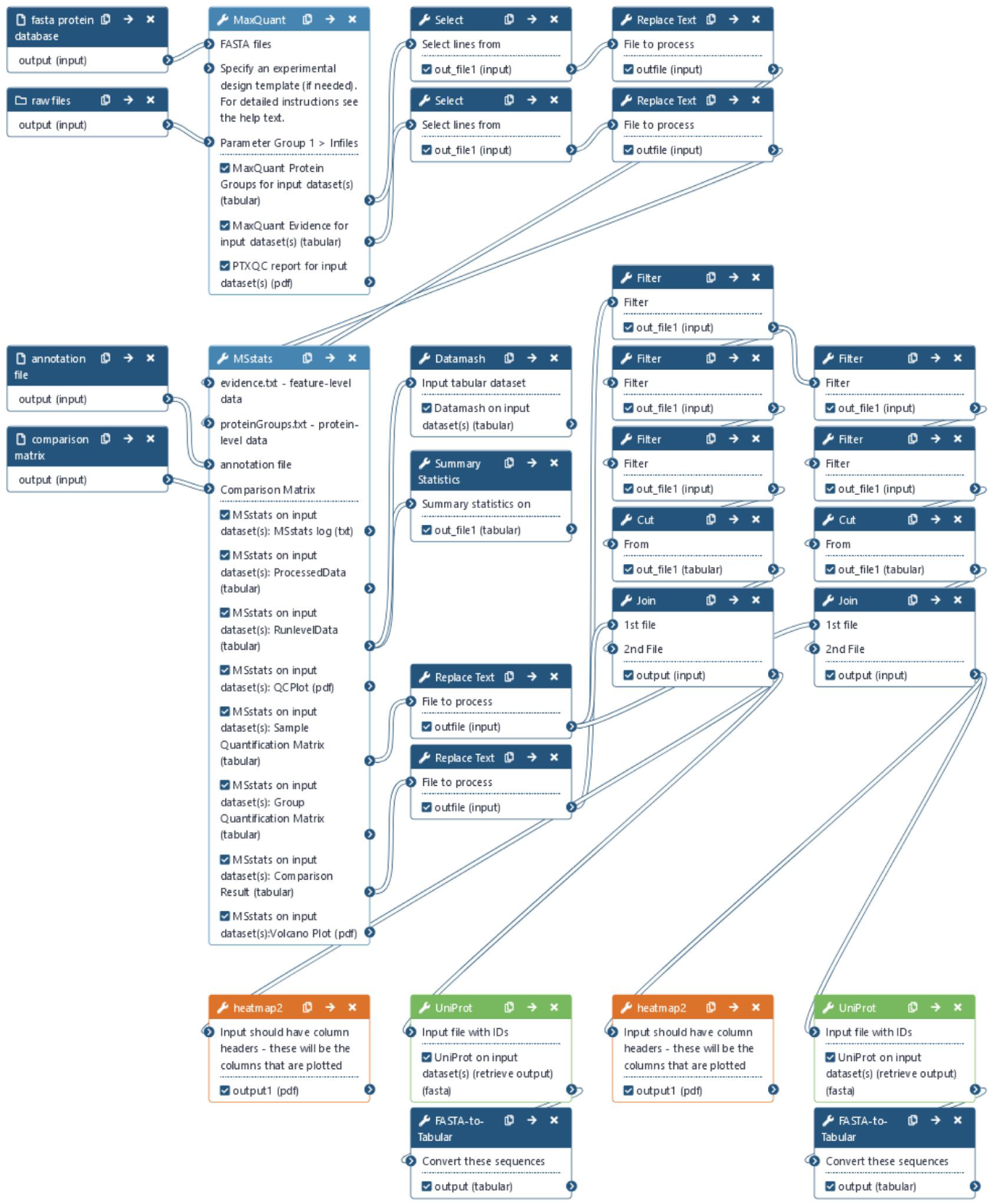
Galaxy provides a workflow management system and high levels of interoperability due to its modular tools. Depicted is a label-free proteomics workflow from a tutorial in the Galaxy training network. The workflow is based on MaxQuant and MSstats (light blue). Both tools are chained together with already existing Galaxy tools for filtering and text manipulation (dark blue), visualization (yellow) and protein annotation (green). Users only need to load the required input files (raw, FASTA, MSstats annotation file and MSstats comparison matrix) into a new analysis history and select them as inputs for the workflow. Then all workflow steps run automatically, however email notification can be enabled when selected tools are finished.

### Training material with example datasets

To facilitate the usage of these newly built quantitative proteomics tools in Galaxy, we created three accompanying tutorials that showcase the application of MaxQuant and MSstats for different use cases. All three trainings are available online via the central repository of the Galaxy Training Network ^21^ (Table S1) and provide example datasets and step-by-step explanations that enable hands-on training.

The first training is tailored towards researchers that are not yet familiar with MaxQuant. Two human serum samples are analyzed. One sample was depleted for the most abundant serum proteins and the training aims to find which of the samples was depleted and how successful the depletion was. To answer this question, a label-free MaxQuant analysis is performed, followed by inspecting the quality control report from the PTXQC tool, filtering, sorting, computing and visualizing the properties of both datasets (Figure 3a).

**Figure 3:**
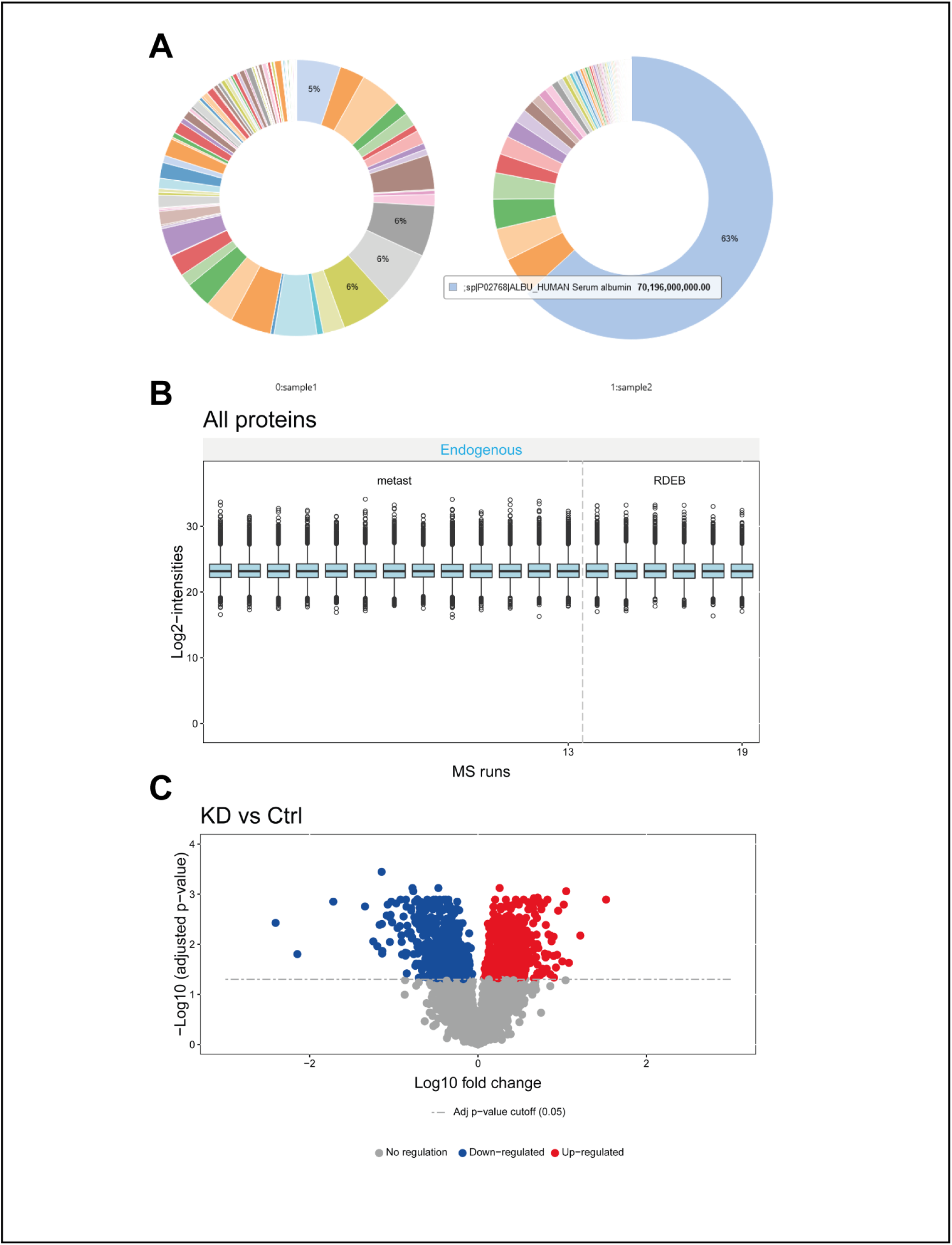
Three distinct trainings based on MaxQuant and MSstats are available in the Galaxy Training Network (GTN): https://training.galaxyproject.org. A) Interactive pie chart that shows the protein abundances of the depleted serum sample (sample 1, on the left) and the protein abundance of the non-depleted serum sample (sample 2, on the right). B) MSstats plot of log2 intensity distributions for 19 patients with two different skin cancers: Metastasizing squamous cell carcinoma (metast) and Recessive Dystrophic Epidermolysis Bullosa squamous cell carcinoma (RDEB). C) MSstatsTMT volcano plot showing the p-values and fold changes for knockdown vs. control cells.

The second training explores a realistic label-free dataset consisting of skin cancer tissue samples from 19 patients ^22^. The training starts with a label-free analysis in MaxQuant, followed by statistical analysis in MSstats to find differentially abundant proteins between two types of skin cancers. Several follow-up steps are performed to filter and visualize the result and annotate proteins of interest (Figure 3b).

The third training explores a realistic, fractionated TMT dataset consisting of 12 high pH fractions from a human cell line experiment ^23^. The training starts with the TMT11-plex analysis in MaxQuant followed by the statistical analysis in MSstatsTMT to find differentially abundant proteins between knockdown and control cells (Figure 3c).

The trainings do not only serve as self-study material but due to the detailed descriptions and the hands-on design, are a meaningful resource to teach proteomics data analysis to researchers and (undergraduate) students in their curriculum ^24^.

## Conclusion

The integration of MaxQuant and MSstats into the Galaxy framework allows easily accessible, reproducible and scalable quantitative proteomics data analysis. An internet connection and web browser suffice to run these tools on public clouds. The availability of many other omics tools in the Galaxy framework allows the integration of MaxQuant and MSstats into more complex, even multi-omics analyses in a single analysis platform. In addition, the Galaxy framework enables the highest levels of reproducible research starting from tool version control to storing all metadata and intermediate results. This enabled MaxQuant in combination with MSstats for the first time to run in an accessible and reproducible way, in parallel on large infrastructures, which is the next step to real high-throughput proteomics.

## Availability

- Galaxy toolshed: https://toolshed.g2.bx.psu.edu
- European Galaxy server: https://usegalaxy.eu
- Galaxy Training Network: https://training.galaxyproject.org/training-material/topics/proteomics
- Github repository: https://github.com/galaxyproteomics/tools-galaxyp

## Acknowledgment

The authors thank the Galaxy community for critically reviewing tools and training materials and Tim Dudgeon for support with the MaxQuant wrapper. The authors thank Olga Vitek, Meena Choi, Ting Huang and Mateusz Staniak (Northeastern University) and Derya Steenbuck (University of Freiburg) for providing test files and for helpful discussions. The authors acknowledge the support of the Freiburg Galaxy Team, Bioinformatics, University of Freiburg (Germany) funded by the Collaborative Research Centre 992 Medical Epigenetics (DFG grant SFB 992/1 2012) and the German Federal Ministry of Education and Research BMBF (grant 031 A538A de.NBI-RBC). OS acknowledges funding by the Deutsche Forschungsgemeinschaft (DFG, SCHI 871/17-1, NY 90/6-1, SCHI 871/15-1, GR 4553/5-1, PA 2807/3-1, project-ID 431984000 - SFB 1453, project-ID 441891347 - SFB 1479, project-ID 423813989 - GRK 2606, project-ID 322977937 - GRK 2344, www.pu), the ERA PerMed programme (BMBF, 01KU1916, 01KU1915A), the German-Israel Foundation (grant no. 1444), and the German Consortium for Translational Cancer Research (project Impro-Rec). BW acknowledges support by the DFG, Project-ID 403222702/SFB 1381, TRR 130 and FOR 2743.

## Supporting information

- Table S1: Overview and links to the Galaxy tools and training materials (XLSX)

## References

(1) Cox, J.; Mann, M. MaxQuant Enables High Peptide Identification Rates, Individualized p.p.b.-Range Mass Accuracies and Proteome-Wide Protein Quantification. Nat. Biotechnol. 2008, 26 (12), 1367–1372. https://doi.org/10.1038/nbt.1511.

(2) Sinitcyn, P.; Tiwary, S.; Rudolph, J.; Gutenbrunner, P.; Wichmann, C.; Yllmaz, Ş.; Hamzeiy, H.; Salinas, F.; Cox, J. MaxQuant Goes Linux. Nat. Methods 2018, 15 (6), 401. https://doi.org/10.1038/s41592-018-0018-y.

(3) Choi, M.; Chang, C.-Y.; Clough, T.; Broudy, D.; Killeen, T.; MacLean, B.; Vitek, O. MSstats: An R Package for Statistical Analysis of Quantitative Mass Spectrometry-Based Proteomic Experiments. Bioinformatics 2014, 30 (17), 2524–2526. https://doi.org/10.1093/bioinformatics/btu305.

(4) Huang, T.; Choi, M.; Tzouros, M.; Golling, S.; Pandya, N. J.; Banfai, B.; Dunkley, T.; Vitek, O. MSstatsTMT: Statistical Detection of Differentially Abundant Proteins in Experiments with Isobaric Labeling and Multiple Mixtures. Mol. Cell. Proteomics 2020, 19 (10), 1706–1723. https://doi.org/10.1074/mcp.RA120.002105.

(5) Cox, J.; Hein, M. Y.; Luber, C. A.; Paron, I.; Nagaraj, N.; Mann, M. Accurate Proteome-Wide Label-Free Quantification by Delayed Normalization and Maximal Peptide Ratio Extraction, Termed MaxLFQ. Mol. Cell. Proteomics 2014, 13 (9), 2513–2526. https://doi.org/10.1074/mcp.M113.031591.

(6) Bielow, C.; Mastrobuoni, G.; Kempa, S. Proteomics Quality Control: Quality Control Software for MaxQuant Results. J. Proteome Res. 2016, 15 (3), 777–787. https://doi.org/10.1021/acs.jproteome.5b00780.

(7) Perez-Riverol, Y.; Moreno, P. Scalable Data Analysis in Proteomics and Metabolomics Using BioContainers and Workflows Engines; Wiley-VCH Verlag, 2020; Vol. 20. https://doi.org/10.1002/pmic.201900147.

(8) Neely, B. A. Cloudy with a Chance of Peptides: Accessibility, Scalability, and Reproducibility with Cloud-Hosted Environments. J. Proteome Res. 2021, acs.jproteome.0c00920. https://doi.org/10.1021/acs.jproteome.0c00920.

(9) Grüning, B.; Chilton, J.; Köster, J.; Dale, R.; Soranzo, N.; van den Beek, M.; Goecks, J.; Backofen, R.; Nekrutenko, A.; Taylor, J. Practical Computational Reproducibility in the Life Sciences. Cell Syst. 2018, 6 (6), 631–635. https://doi.org/10.1016/j.cels.2018.03.014.

(10) Afgan, E.; Baker, D.; Batut, B.; van den Beek, M.; Bouvier, D.; Čech, M.; Chilton, J.; Clements, D.; Coraor, N.; Grüning, B. A.; Guerler, A.; Hillman-Jackson, J.; Hiltemann, S.; Jalili, V.; Rasche, H.; Soranzo, N.; Goecks, J.; Taylor, J.; Nekrutenko, A.; Blankenberg, D. The Galaxy Platform for Accessible, Reproducible and Collaborative Biomedical Analyses: 2018 Update. Nucleic Acids Res. 2018, 46 (W1), W537–W544. https://doi.org/10.1093/nar/gky379.

(11) Chambers, M. C.; Maclean, B.; Burke, R.; Amodei, D.; Ruderman, D. L.; Neumann, S.; Gatto, L.; Fischer, B.; Pratt, B.; Egertson, J.; Hoff, K.; Kessner, D.; Tasman, N.; Shulman, N.; Frewen, B.; Baker, T. A.; Brusniak, M.-Y.; Paulse, C.; Creasy, D.; Flashner, L.; Kani, K.; Moulding, C.; Seymour, S. L.; Nuwaysir, L. M.; Lefebvre, B.; Kuhlmann, F.; Roark, J.; Rainer, P.; Detlev, S.; Hemenway, T.; Huhmer, A.; Langridge, J.; Connolly, B.; Chadick, T.; Holly, K.; Eckels, J.; Deutsch, E. W.; Moritz, R. L.; Katz, J. E.; Agus, D. B.; MacCoss, M.; Tabb, D. L.; Mallick, P. A Cross-Platform Toolkit for Mass Spectrometry and Proteomics. Nat. Biotechnol. 2012, 30 (10), 918–920. https://doi.org/10.1038/nbt.2377.

(12) Barsnes, H.; Vaudel, M. SearchGUI: A Highly Adaptable Common Interface for Proteomics Search and de Novo Engines. J. Proteome Res. 2018, 17 (7), 2552–2555. https://doi.org/10.1021/acs.jproteome.8b00175.

(13) Vaudel, M.; Burkhart, J. M.; Zahedi, R. P.; Oveland, E.; Berven, F. S.; Sickmann, A.; Martens, L.; Barsnes, H. PeptideShaker Enables Reanalysis of MS-Derived Proteomics Data Sets. Nat. Biotechnol. 2015, 33 (1), 22–24. https://doi.org/10.1038/nbt.3109.

(14) Röst, H. L.; Sachsenberg, T.; Aiche, S.; Bielow, C.; Weisser, H.; Aicheler, F.; Andreotti, S.; Ehrlich, H.-C.; Gutenbrunner, P.; Kenar, E.; Liang, X.; Nahnsen, S.; Nilse, L.; Pfeuffer, J.; Rosenberger, G.; Rurik, M.; Schmitt, U.; Veit, J.; Walzer, M.; Wojnar, D.; Wolski, W. E.; Schilling, O.; Choudhary, J. S.; Malmström, L.; Aebersold, R.; Reinert, K.; Kohlbacher, O. OpenMS: A Flexible Open-Source Software Platform for Mass Spectrometry Data Analysis. Nat. Methods 2016, 13 (9), 741–748. https://doi.org/10.1038/nmeth.3959.

(15) Röst, H. L.; Rosenberger, G.; Navarro, P.; Gillet, L.; Miladinović, S. M.; Schubert, O. T.; Wolski, W.; Collins, B. C.; Malmström, J.; Malmström, L.; Aebersold, R. OpenSWATH Enables Automated, Targeted Analysis of Data-Independent Acquisition MS Data. Nat. Biotechnol. 2014, 32 (3), 219–223. https://doi.org/10.1038/nbt.2841.

(16) Tsou, C. C.; Avtonomov, D.; Larsen, B.; Tucholska, M.; Choi, H.; Gingras, A. C.; Nesvizhskii, A. I. DIA-Umpire: Comprehensive Computational Framework for Data-Independent Acquisition Proteomics. Nat. Methods 2015, 12 (3), 258–264. https://doi.org/10.1038/nmeth.3255.

(17) Grüning, B.; Dale, R.; Sjödin, A.; Chapman, B. A.; Rowe, J.; Tomkins-Tinch, C. H.; Valieris, R.; Köster, J. Bioconda: Sustainable and Comprehensive Software Distribution for the Life Sciences. Nat. Methods 2018, 15 (7), 475–476. https://doi.org/10.1038/s41592-018-0046-7.

(18) da Veiga Leprevost, F.; Grüning, B. A.; Alves Aflitos, S.; Röst, H. L.; Uszkoreit, J.; Barsnes, H.; Vaudel, M.; Moreno, P.; Gatto, L.; Weber, J.; Bai, M.; Jimenez, R. C.; Sachsenberg, T.; Pfeuffer, J.; Vera Alvarez, R.; Griss, J.; Nesvizhskii, A. I.; Perez-Riverol, Y. BioContainers: An Open-Source and Community-Driven Framework for Software Standardization. Bioinformatics 2017, 33 (16), 2580–2582. https://doi.org/10.1093/bioinformatics/btx192.

(19) Pedrioli, P. G. A.; Eng, J. K.; Hubley, R.; Vogelzang, M.; Deutsch, E. W.; Raught, B.; Pratt, B.; Nilsson, E.; Angeletti, R. H.; Apweiler, R.; Cheung, K.; Costello, C. E.; Hermjakob, H.; Huang, S.; Julian, R. K.; Kapp, E.; McComb, M. E.; Oliver, S. G.; Omenn, G.; Paton, N. W.; Simpson, R.; Smith, R.; Taylor, C. F.; Zhu, W.; Aebersold, R. A Common Open Representation of Mass Spectrometry Data and Its Application to Proteomics Research; Nat Biotechnol, 2004; Vol. 22. https://doi.org/10.1038/nbt1031.

(20) Martens, L.; Chambers, M.; Sturm, M.; Kessner, D.; Levander, F.; Shofstahl, J.; Tang, W. H.; Rompp, A.; Neumann, S.; Pizarro, A. D.; Montecchi-Palazzi, L.; Tasman, N.; Coleman, M.; Reisinger, F.; Souda, P.; Hermjakob, H.; Binz, P.-A.; Deutsch, E. W. MzML-a Community Standard for Mass Spectrometry Data. Mol. Cell. Proteomics 2011, 10 (1), R110.000133–R110.000133. https://doi.org/10.1074/mcp.R110.000133.

(21) Batut, B.; Hiltemann, S.; Bagnacani, A.; Baker, D.; Bhardwaj, V.; Blank, C.; Bretaudeau, A.; Brillet-Guéguen, L.; Čech, M.; Chilton, J.; Clements, D.; Doppelt-Azeroual, O.; Erxleben, A.; Freeberg, M. A.; Gladman, S.; Hoogstrate, Y.; Hotz, H.-R.; Houwaart, T.; Jagtap, P.; Larivière, D.; Le Corguillé, G.; Manke, T.; Mareuil, F.; Ramírez, F.; Ryan, D.; Sigloch, F. C.; Soranzo, N.; Wolff, J.; Videm, P.; Wolfien, M.; Wubuli, A.; Yusuf, D.; Taylor, J.; Backofen, R.; Nekrutenko, A.; Grüning, B. Community-Driven Data Analysis Training for Biology. Cell Syst. 2018, 6 (6), 752–758.e1. https://doi.org/10.1016/j.cels.2018.05.012.

(22) Föll, M. C.; Fahrner, M.; Gretzmeier, C.; Thoma, K.; Biniossek, M. L.; Kiritsi, D.; Meiss, F.; Schilling, O.; Nyström, A.; Kern, J. S. Identification of Tissue Damage, Extracellular Matrix Remodeling and Bacterial Challenge as Common Mechanisms Associated with High-Risk Cutaneous Squamous Cell Carcinomas. Matrix Biol. 2018, 66, 1–21. https://doi.org/10.1016/j.matbio.2017.11.004.

(23) Baumert, H. M.; Metzger, E.; Fahrner, M.; George, J.; Thomas, R. K.; Schilling, O.; Schüle, R. Depletion of Histone Methyltransferase KMT9 Inhibits Lung Cancer Cell Proliferation by Inducing Non-Apoptotic Cell Death. Cancer Cell Int. 2020, 20 (1), 52. https://doi.org/10.1186/s12935-020-1141-2.

(24) Serrano-Solano, B.; Föll, M. C.; Gallardo-Alba, C.; Erxleben, A.; Rasche, H.; Hiltemann, S.; Fahrner, M.; Dunning, M. J.; Schulz, M. H.; Scholtz, B.; Clements, D.; Nekrutenko, A.; Batut, B.; Grüning, B. A. Fostering Accessible Online Education Using Galaxy as an E-Learning Platform. PLOS Comput. Biol. 2021, 17 (5), e1008923. https://doi.org/10.1371/journal.pcbi.1008923.

(25) Blankenberg, D.; Von Kuster, G.; Bouvier, E.; Baker, D.; Afgan, E.; Stoler, N.; Taylor, J.; Nekrutenko, A.; Clements, D.; Coraor, N.; Eberhard, C.; Francheteau, D.; Goecks, J.; Guerler, S.; Jackson, J.; Cooke, I.; Johnson, J.; Kirton, E.; Cock, P.; Chapman, B.; Grüning, B.; Lazarus, R. Dissemination of Scientific Software with Galaxy ToolShed. Genome Biol. 2014, 15 (2), 403. https://doi.org/10.1186/gb4161.

(26) Mehta, S.; Easterly, C. W.; Sajulga, R.; Millikin, R. J.; Argentini, A.; Eguinoa, I.; Martens, L.; Shortreed, M. R.; Smith, L. M.; McGowan, T.; Kumar, P.; Johnson, J. E.; Griffin, T. J.; Jagtap, P. D. Precursor Intensity-Based Label-Free Quantification Software Tools for Proteomic and Multi-Omic Analysis within the Galaxy Platform. Proteomes 2020, 8 (3), 15. https://doi.org/10.3390/proteomes8030015.

(27) Stewart, P. A.; Kuenzi, B. M.; Mehta, S.; Kumar, P.; Johnson, J. E.; Jagtap, P.; Griffin, T. J.; Haura, E. B. The Galaxy Platform for Reproducible Affinity Proteomic Mass Spectrometry Data Analysis. In Mass Spectrometry of Proteins; Evans, C. A., Wright, P. C., Noirel, J., Eds.; Methods in Molecular Biology; Springer New York: New York, NY, 2019; Vol. 1977, pp 249–261. https://doi.org/10.1007/978-1-4939-9232-4_16.

(28) Jagtap, P. D.; Johnson, J. E.; Onsongo, G.; Sadler, F. W.; Murray, K.; Wang, Y.; Shenykman, G. M.; Bandhakavi, S.; Smith, L. M.; Griffin, T. J. Flexible and Accessible Workflows for Improved Proteogenomic Analysis Using the Galaxy Framework. J. Proteome Res. 2014, 13 (12), 5898–5908. https://doi.org/10.1021/pr500812t.

(29) Davidson, R. L.; Weber, R. J. M.; Liu, H.; Sharma-Oates, A.; Viant, M. R. Galaxy-M: A Galaxy Workflow for Processing and Analyzing Direct Infusion and Liquid Chromatography Mass Spectrometry-Based Metabolomics Data. GigaScience 2016, 5 (1), 10. https://doi.org/10.1186/s13742-016-0115-8.

(30) Peters, K.; Bradbury, J.; Bergmann, S.; Capuccini, M.; Cascante, M.; de Atauri, P.; Ebbels, T. M. D.; Foguet, C.; Glen, R.; Gonzalez-Beltran, A.; Günther, U. L.; Handakas, E.; Hankemeier, T.; Haug, K.; Herman, S.; Holub, P.; Izzo, M.; Jacob, D.; Johnson, D.; Jourdan, F.; Kale, N.; Karaman, I.; Khalili, B.; Emami Khonsari, P.; Kultima, K.; Lampa, S.; Larsson, A.; Ludwig, C.; Moreno, P.; Neumann, S.; Novella, J. A.; O’Donovan, C.; Pearce, J. T. M.; Peluso, A.; Piras, M. E.; Pireddu, L.; Reed, M. A. C.; Rocca-Serra, P.; Roger, P.; Rosato, A.; Rueedi, R.; Ruttkies, C.; Sadawi, N.; Salek, R. M.; Sansone, S. A.; Selivanov, V.; Spjuth, O.; Schober, D.; Thévenot, E. A.; Tomasoni, M.; van Rijswijk, M.; van Vliet, M.; Viant, M. R.; Weber, R. J. M.; Zanetti, G.; Steinbeck, C. PhenoMeNal: Processing and Analysis of Metabolomics Data in the Cloud. GigaScience 2019, 8 (2), 1–12. https://doi.org/10.1093/gigascience/giy149.

(31) Guitton, Y.; Tremblay-Franco, M.; Le Corguillé, G.; Martin, J.-F.; Pétéra, M.; Roger-Mele, P.; Delabrière, A.; Goulitquer, S.; Monsoor, M.; Duperier, C.; Canlet, C.; Servien, R.; Tardivel, P.; Caron, C.; Giacomoni, F.; Thévenot, E. A. Create, Run, Share, Publish, and Reference Your LC-MS, FIA-MS, GC-MS, and NMR Data Analysis Workflows with the Workflow4Metabolomics 3.0 Galaxy Online Infrastructure for Metabolomics. Int. J. Biochem. Cell Biol. 2017, 93, 89–101. https://doi.org/10.1016/j.biocel.2017.07.002.

(32) Boekel, J.; Chilton, J. M.; Cooke, I. R.; Horvatovich, P. L.; Jagtap, P. D.; Käll, L.; Lehtiö, J.; Lukasse, P.; Moerland, P. D.; Griffin, T. J. Multi-Omic Data Analysis Using Galaxy. Nat. Biotechnol. 2015, 33 (2), 137–139. https://doi.org/10.1038/nbt.3134.

(33) Fahrner, M.; Föll, M. C.; Grüning, B.; Bernt, M.; Röst, H.; Schilling, O. Democratizing Data-Independent Acquisition Proteomics Analysis on Public Cloud Infrastructures Via The Galaxy Framework; preprint; bioRxiv, 2021. https://doi.org/10.1101/2021.07.21.453197.

